# How Sensitive are EEG Results to Preprocessing Methods: A Benchmarking Study

**DOI:** 10.1101/2020.01.20.913327

**Authors:** Kay A. Robbins, Jonathan Touryan, Tim Mullen, Christian Kothe, Nima Bigdely-Shamlo

## Abstract

Although several guidelines for best practices in EEG preprocessing have been released, even those studies that strictly adhere to those guidelines contain considerable variation in the ways that the recommended methods are applied. An open question for researchers is how sensitive the results of EEG analyses are to variations in preprocessing methods and parameters. To address this issue, we analyze the effect of preprocessing methods on downstream EEG analysis using several simple signal and event-related measures. Signal measures include recording-level channel amplitudes, study-level channel amplitude dispersion, and recording spectral characteristics. Event-related methods include ERPs and ERSPs and their correlations across methods for a diverse set of stimulus events. Our analysis also assesses differences in residual signals both in the time and spectral domains after blink artifacts have been removed. Using fully automated pipelines, we evaluate these measures across 17 EEG studies for two ICA-based preprocessing approaches (LARG, MARA) plus two variations of Artifact Subspace Reconstruction (ASR). Although the general structure of the results is similar across these preprocessing methods, there are significant differences, particularly in the low-frequency spectral features and in the residuals left by blinks. These results argue for detailed reporting of processing details as suggested by most guidelines, but also for using a federation of automated processing pipelines and comparison tools to quantify effects of processing choices as part of the research reporting.

## I. Introduction

EEG (electroencephalography) is widely used to record brain activity in clinical, research laboratory, and real-world settings. Although a number of guidelines for best practices in processing EEG have appeared in recent years (see for example, [1] [2] [3]), the guidelines are quite broad and give researchers significant leeway in creating compliant processing pipelines. A crucial question for evaluating the reliability and comparability of results from different studies is how details of the processing pipelines might influence the end results.

This paper begins to address the preprocessing variability question by assessing differences in signal distributions across studies for several different preprocessing methods. We also examine differences in event-related potentials and event-related spectral perturbations computed by both trial averaging (ERPs and ERSPs, respectively) and temporal overlap regression (rERPs and rERSPs, respectively) [4] [5] [6]. The remainder of the paper is organized as follows. Section II describes briefly describes the data corpus, delineates the four pipelines compared in this paper, and introduces the signal and event-related feature metrics used for evaluation. Section III presents comparisons organized by signal and event-related feature characteristics, with special evaluation of the effect of blinks. Section IV discusses some of the implications of the results, and Section V gives some concluding remarks.

## II. Methods and materials

The data corpus for this study consists of EEG recordings from 17 studies performed at six experimental sites containing approximately 7.8 million event-related epochs from 1,100 recordings as described [7]. The studies fall into two general categories: visual target detection and lane-keeping tasks that include distractions and other variations. The raw data and some levels of processed data are available through the DataCatalog at https://cancta.net. Code for the preprocessing pipelines and calculation of some of the metrics is available at https://github.com/VisLab/EEG-Pipelines. The study events were annotated using Hierarchical Event descriptors (HED tags) prior to any processing [8].

### A. Early-stage preprocessing

We applied the PREP pipeline [9] to remove line noise, identify bad channels, and robust average reference the data. If bad channels were interpolated, EEGLAB’s *eeg_interp()* was used for spherical interpolation, and re-sampling was done with EEGLAB’s *pop_resample()*. All filtering operations used EEGLAB’s *pop_eegfiltnew()* with the default settings unless otherwise specified. Datasets with more than 64 channels were reduced to the 64 channels closest to the standard 10-20 positions and assigned standard 10-20 labels [7]. Blink events, identified as the positions of the maximum amplitude of the blink, were then inserted into the EEG structure using the automated BLINKER toolbox [10].

### B. Preprocessing pipelines used in this study

This work compares four processing pipelines, denoted as LARG, MARA, ASR_5* and ASR_10*, respectively. The LARG [7] and MARA [11] are closely-related ICA-based pipelines, while the ASR_x* pipelines are based on the Artifact Subspace Reconstruction algorithm [12], an automated EEG artifact removal algorithm that can be applied in real-time. The full ASR pipeline and is now part of the recommended preprocessing pipeline for EEGLAB [13] [14]. (See Suppl. Fig. 1 for a summary diagram of the pipelines.)

#### 1) The LARG pipeline

LARG [7] is an automated pipeline that emphasizes the removal of eye artifacts. After interpolating the bad channels identified by PREP, LARG uses the default settings of EEGLAB *pop_eegfiltnew()* to high-pass filter the data at 1 Hz with a zero-phase FIR filter and a Hamming window. After down-sampling to 128 Hz, LARG computes independent components (ICs) using Infomax applied to cleaned sections of the data as described in [15]. Some studies were processed using the CUDAICA GPU implementation of Infomax [16]. LARG removes from the signal ICs identified by EyeCatch [17] as eye artifacts, and applies temporal overlap regression to remove the residual time-domain blinks in intervals of [−1, 1].

#### 2) The MARA pipeline

MARA (Multiple Artifact Rejection Algorithm) automates the selection of artifactual independent components ICs by applying multiple statistical tests [18]. Our MARA pipeline uses a pipeline identical to LARG except that instead of applying EyeCatch and regressing out blinks, the MARA pipeline removes artifactual ICs based on the MARA criteria.

#### 3) The ASR* pipeline (ASR_5* and ASR_10*)

The ASR (Artifact Subspace Reconstruction) algorithm [12] uses principal-component-like subspace decomposition to eliminate large transients. ASR can be applied in an online setting for real-time artifact removal. We applied ASR using the *clean_asr()* function from the *clean_rawdata* EEGLAB plugin. Note that the recommended ASR artifact removal pipeline and the default approach implemented in *clean_rawdata* includes bad channel removal and bad window removal, which can significantly improve artifact removal as well as its own filtering. However, here we performed comparisons with only underlying subspace removal implemented in the *clean_asr()* function, without the benefits of these additional offline artifact removal steps. We therefore denote this approach as ASR*. We use the non-interpolated average referenced signal produced by PREP, remove the channel means, and high-pass filter using the default settings for *pop_eegfiltnew()* with a cutoff of 1.5 Hz. The higher cutoff (compared to a 1 Hz cutoff for LARG and MARA) was needed to achieve suitable stop-band suppression below 0.5 Hz for some recordings that had significant drift thereby ensuring the data had a mean of approximately zero within short windows (an essential stationarity pre-condition for ASR to function properly).

ASR has a configurable burst cutoff parameter for determining how aggressively it removes transient high-variance artifacts, with smaller values corresponding to more aggressive removal of artifacts. We used burst cutoff parameters: 5 (highly aggressive) and 10 (modestly aggressive, typical setting), denoting the pipelines as ASR_5* and ASR_10*, respectively. ASR calibration was performed separately for each recording using the default settings, which identify a signal subspace from clean segments of the entire recording after removal of segments of data containing high-power artifacts in more than 7.5% of channels.

### C. Computation of robust channel signal statistics

Bigdely-Shamlo et al. [19] introduced several robust summary metrics for signal channel distributions which we use here, including the *recording channel amplitude vector* and the *study channel dispersion vector*. These metrics capture the signal scale across channels in a recording, and the dispersion of that scale across a study, respectively. Due to the robust estimators being used, these measures are partially biased towards brain signals rather than artifacts, and can thus be used to track impacts on those brain signals before and after a given pre-processing method is applied.

Prior to calculation of the summary features, the EEG signal is filtered using a [1, 20] Hz bandpass FIR filter and the following 10-20 standard set of 26 channels is selected: FP1, FP2, F3, Fz, F4, F7, F8, FC3, FCz, FC4, FT7, FT8, C3, Cz, C4, TP7, TP8, CP3, CPz, CP4, P3, Pz, P4, O1, Oz, and O2. All recordings in the corpus contain these 26 channels, which are used for the channel summary metrics unless otherwise stated.

The *recording channel amplitude vector* is a 26×1 positive vector of the robust standard deviations (defined as 1.4826 × the median absolute deviation from the sample median) of the filtered channel signals from these 26 common channels. The *study amplitude matrix* is a 26×*S* matrix of the recording channel amplitude vectors stacked across the *S* recordings in the study. The *corpus amplitude matrix A* is a 26×*C* matrix formed by stacking the study amplitude matrices across all of the studies in the corpus. Here *C* is the total number of recordings in the corpus. The *dispersion vector* for a study or corpus amplitude matrix is a 26×1 positive vector calculated as the robust standard deviation of each row of the respective amplitude matrix divided by the median of that row. (See [7] for a more detailed description of these metrics.)

Bigdely-Shamlo et al. [7] also showed that dividing the recording channel data by a recording-specific constant prior to computing the study or corpus dispersion vector greatly reduces the dispersion values. Several methods of computing the recording-specific constant were shown to be effective in reducing study-wide channel dispersion. Here we use the Huber mean of the recording channel amplitude vector as the recording-specific constant. The *normalized study amplitude matrix* and the *normalized corpus amplitude matrix* in this paper are formed by dividing each column of the respective amplitude matrix by its column Huber mean.

To visualize, we take the median of the rows of the corpus amplitude matrix and plot the resulting 26×1 vector using EEGLAB’s *topoplot()*. To explore channel signal dependencies on recording-specific scaling, we plotted entry A(*i, k*) versus A(*j, k*) (with *i ≠ j*). A ball-shaped plot indicates that little of the recording variability can be addressed by this recording-specific normalization, while a linear shape suggests that such normalization will improve comparability.

To quantify to what extent dividing each recording by a recording-specific constant reduces channel dispersion across a corpus, we calculated the percentage of dispersion reduction for each study, channel, and method separately using the formula 100*(*dispersion before* – *dispersion after*)/(*dispersion before*). We then averaged these percentages for each preprocessing method to obtain an overall dispersion reduction percentage [7].

### D. Computation of channel spectral characteristics

To see how different preprocessing approaches might distort the signal spectral characteristics, we calculated both summary and local measures as follows. Each recording was scaled by a recording-specific constant (the Huber mean of the recording amplitude vector). We computed the time-varying spectral decomposition of each of the 26 common channels by applying the MATLAB continuous wavelet transform *cwt()* using the complex Morlet wavelet family *cmor*1-1.5 and 50 frequencies logarithmically sampled in the range 2 to 30 Hz. We then normalized the amplitude at each frequency for each spectrogram by subtracting the median over time and dividing by the median absolute deviation from the median (MAD). We refer to this operation as robust z-scoring.

For each preprocessing method, we created a *spectral fingerprint* of each recording by vectorizing the normalized spectrograms. We then computed correlations of the corresponding fingerprint vectors associated with pairs of preprocessing methods to summarize how much preprocessing affects spectral results. In addition, we averaged each spectrogram within standard frequency bands (delta: [2, 4] Hz, theta: [4, 7] Hz, alpha: [7, 12] Hz, beta: [12, 30] Hz) to form separate fingerprints for each band and computed band correlations for pairs of preprocessing methods.

For each preprocessing method, we also created a *recording spectral sample* by choosing at random 100 non-overlapping segments of 4 seconds duration from each recording and calculating the power spectral density (PSD) of each sample segment. We used the Matlab *pmtm()* multi-taper spectral density function with tapers having a half-bandwidth of 4 using 512 points and 256 frequency bins in [1, 50] Hz. PSD samples were normalized by dividing by total spectral power, similar to those used by Cruz-Garza et al. [20]. We then computed the Pearson correlation (across frequency) between PSD samples for different pairs of preprocessing methods. This metric quantifies the relationship for each recording by 26×100 = 2,600 correlations rather than via a single correlation value.

We also computed the mean spectra for each spectral sample in each of five specified frequency bands (the delta, theta, alpha, and beta bands listed above, as well as a gamma band of [30, 50] Hz) for each channel in each recording. We then calculated Pearson correlations between corresponding band spectral samples for pairs of preprocessing methods.

### E. Computation of event-related features

We computed the event-related features on intervals of [−2, 2] seconds time-locked around individual events separately for each preprocessing method. As described in [21], we used two different computation methods: ordinary trial averaging (ERPs and ERSPs) and temporal overlap regression (rERPs and rERSPs). We computed (r)ERPs for each (*recording*, *study-specific event code*, *channel*) and (r)ERSPs for each (*recording*, *study-specific event code*, *channel, frequency*) combination. The (r)ERSPs were computed based on the time-varying amplitude spectrogram computed by applying the MATLAB continuous wavelet transform function, *cwt()*, to the continuous signal at 50 frequencies logarithmically sampled between 2 and 40 Hz. We scaled the resulting amplitudes by subtracting the median and then dividing by 1.4826 times the median, with median computed separately at each frequency over all time points for each recording. We used the outlier detection scheme described in [21] to more robustly compute these features.

Our corpus events were tagged using Hierarchical Event Descriptors (HED tags) to enable cross study comparison. Because many events in our corpus mark non-neurological phenomena such as experimental control, we only considered event codes tagged with *Event/Category/Experimental stimulus* and not also tagged with *Attribute/Offset* for the summary measures. Event-related features corresponding to a particular study-specific event code were only computed for recordings containing at least 10 occurrences of the event code. Event codes that frequently coincided with other event codes were detected and duplicates eliminated. We only considered combinations for which there were at least 5 recordings containing enough events with that event code.

For each of the 26 common channels, we computed the pairwise Pearson correlations, between pairs of preprocessing methods, of the corresponding event-related (r)ERP features (*recording*, *study-specific event code*, *channel*). For (r)ERSP features, we vectorized the spectrograms before computing pairwise correlations. We displayed the resulting distributions of these correlations using boxplots and also performed statistical tests to determine which pairs of preprocessing methods produced more similar event-related features.

### F. Evaluating the effect of blinks

We used the *blink amplitude ratio* to characterize the effect of blink removal for different preprocessing methods. Blink (r)ERPs were computed by time-locking to the *maxFrame* event inserted by BLINKER at the blink amplitude maxima in the EEG signal. We only consider the 26 common channels specified in the previous section. For each (*recording*, *channel*), we baselined the blink (r)ERP by subtracting the mean of the (r)ERP in the time intervals [−2, −1.5] and [1.5, 2] from the entire (r)ERP. We then computed the blink amplitude ratio by dividing the mean absolute value of the baselined blink (r)ERP in the time interval [−0.5, 0.5] by the mean absolute value of the baselined blink signal in the union of the intervals [−2, −1.5] and [1.5, 2]. Ratios close to 1 indicate that the blink signal has been removed during preprocessing without impacting the underlying activity. Ratios much greater than 1 indicate that the blink amplitude has not been fully subtracted from the signal, while ratios close to zero indicate that both the blink and underlying activity have been removed.

## III. Results

### A. Effects of preprocessing on EEG channel statistics

Fig. 1 compares EEG signal properties using the corpus robust channel amplitude matrix, *A.* The top row shows results for data that has been average referenced with bad channels interpolated. The remaining rows correspond to data that has been processed by the LARG, MARA, ASR_10* and ASR_5*, respectively. All signals have been filtered in the range [1, 20] Hz prior to calculation of *A*.

**Fig. 1.**
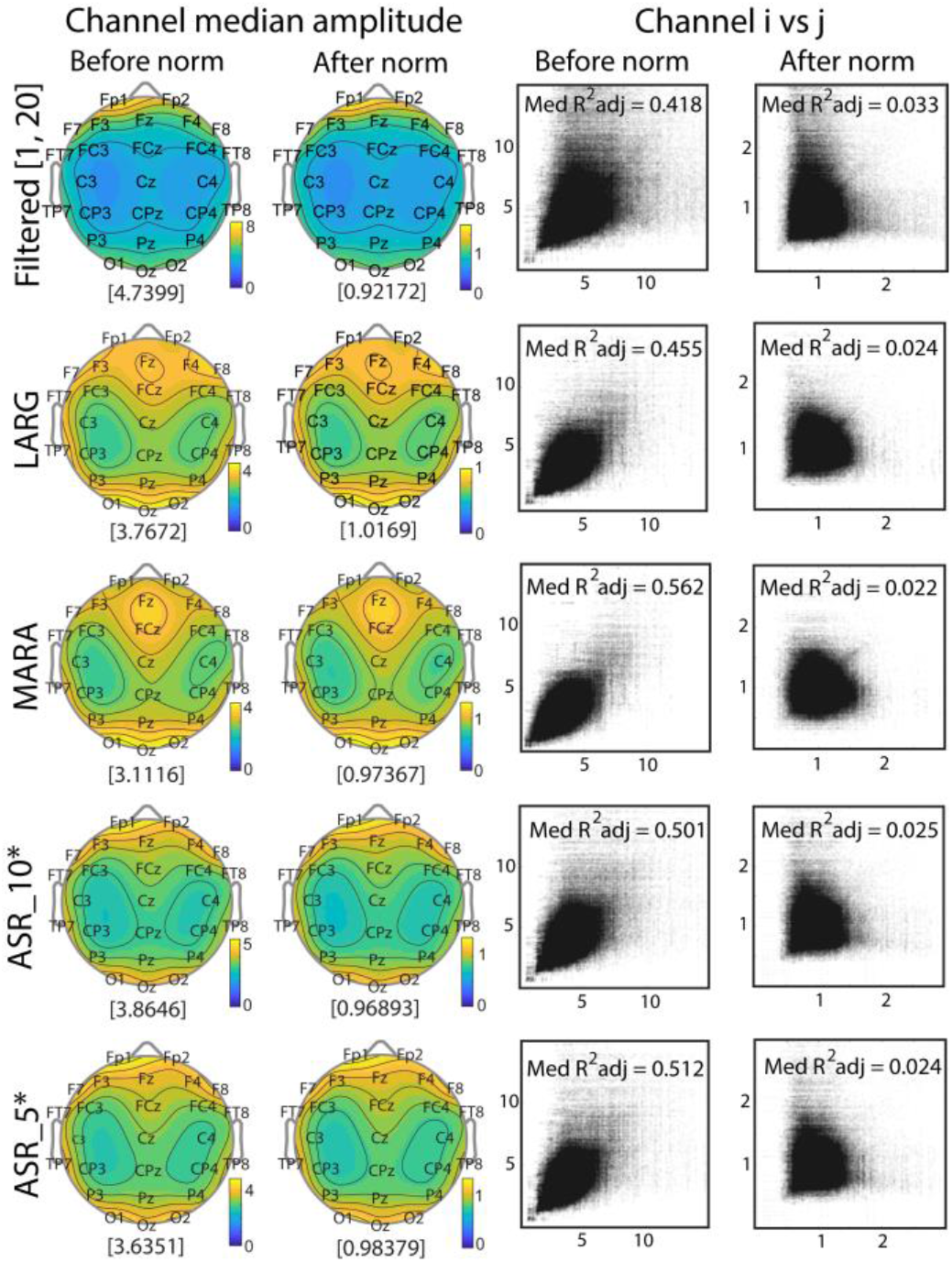
Comparison of summary EEG signal distributions for different preprocessing methods using the 26 common channels. Rows from top to bottom correspond to preprocessing methods: average reference with bad channel interpolation, LARG, MARA, ASR_10*, and ASR_5*. The first column displays scalp maps of the row medians of the corpus amplitude matrix, *A*, while the second column displays scalp maps of the row medians of *A* after normalization by the recording-specific Huber means. Values in square brackets are the medians of the row medians of *A*. The third and fourth columns plot entries *A*(*i*, *k*) versus *A*(*j*, *k*) for all channels *i* ≠ *j* before and after Huber normalization, respectively.

The first column of Fig. 1 shows the row medians of the corpus channel amplitude matrix, *A*, for various processing methods displayed as scalp maps. The scalp maps show a lateral symmetry with a lobe-like structure. Before artifact removal (top row), the signal distributions are dominated by frontal channels due to blinks and other eye artifacts, with additional stronger amplitudes in the occipital regions.

After artifacts have been removed (rows 2 through 5), regardless of processing approach, channel amplitude becomes more equalized across the scalp, with the distinct bilateral lobes becoming more prominent. ASR_10* resembles the average referenced signal the most closely followed by ASR_5*, LARG, and MARA. Both LARG and MARA use ICA-based methods, with MARA removing ICs more aggressively. MARA and LARG show a local maximum near channels Fz and FCz not visible in the ASR variants. We note again, however, that LARG and MARA pipelines applied bad channel removal and interpolation while the ASR* pipelines did not.

The scalp maps after normalization (second column of Fig. 1) have a similar appearance to those prior to normalization, but with a much lower amplitude because normalizing by a constant results in a relative reweighting of the points contributing to the median, keeping the points in a roughly similar relationship.

To investigate whether there is a linear relationship between robust channel amplitudes across recordings (third column of Fig. 1), we plot *A*(*i*, *k*) versus *A*(*j*, *k*) with channel *i* ≠ channel *j* for all recordings *k*. The plots of column 3 show a distinct linear trend irrespective of processing method, indicating the presence of an underlying co-varying relationship. However, the average referenced only data (top row) have many more points on the outer arms, corresponding to the presence of large amplitude blinks and other eye artifacts. The plots corresponding to the other preprocessing methods have much smaller distributions along the axes.

After dividing the channel data by the recording-specific Huber mean normalization factor (an overall robust measure of the recording’s channel amplitude), the *A*(*i*, *k*) versus *A*(*j*, *k*) plots become much less elongated (fourth column of Fig. 1). The top graph of column 4 still has arms, reflecting the continued amplitude dominance of the frontal channels after normalization, as do the ASR variants. The linear channel *i vs j* dependence is greatly reduced as indicated by the median adjusted *R*^2^ values, which are around 0.5 before normalization and nearly 0 afterwards. To quantify the statistical significance of these patterns, we fit a linear regression model to *A*(*i*, *k*) and *A*(*j*, *k*) for each (*i*, *j*) channel pair with *i* ≠ *j*. Table 1 shows the results of this analysis.

**TABLE 1.** Channel *i* vs Channel *j*\*

| Metric<sup>a</sup> | Processing method |  |  |  |  |
| --- | --- | --- | --- | --- | --- |
|  | Filtered | LARG | MARA | ASR_10* | ASR_5* |
| Median adj-$R^2$ (B) | 0.465 | 0.481 | 0.571 | 0.534 | 0.545 |
| Median adj-$R^2$ (A) | 0.023 | 0.015 | 0.015 | 0.018 | 0.018 |
| % sig slopes (B) | 100 | 100 | 100 | 100 | 100 |
| % sig slopes (A) | 76 | 65 | 67 | 73 | 72 |
| # sig channels (B) | 26 | 26 | 23 | 26 | 26 |
| # sig channels (A) | 26 | 26 | 24 | 26 | 26 |

<sup>a</sup> (B) = before normalization, (A) = after normalization.

Before normalization, almost 100% of these 650 linear fits have nonzero slope (*p* < 0.01, *FDR* corrected). The fraction of significant non-zero slopes is reduced to between 0.65 and 0.76 depending on the preprocessing method after normalization. Normalization not only reduces the number of non-zero slopes, but also sharply reduces the quality of the linear fit. This linear relationship, which explains about half of the variability in channel pair amplitudes, almost fully disappears after Huber mean normalization.

Fig. 2 shows that channel dispersion (top graph) is substantially reduced after dividing each recording by its recording-specific Huber mean (bottom graph). The overall average percentage dispersion reduction resulting from dividing each recording by a recording-specific constant ranged from 38% to 45% across studies with no obvious dependence on preprocessing method. The percent reduction was greater than zero with significance *p* < 0.001 (*t*-test, FDR corrected), indicating normalization reduces cross-recording variability

**Fig. 2.**
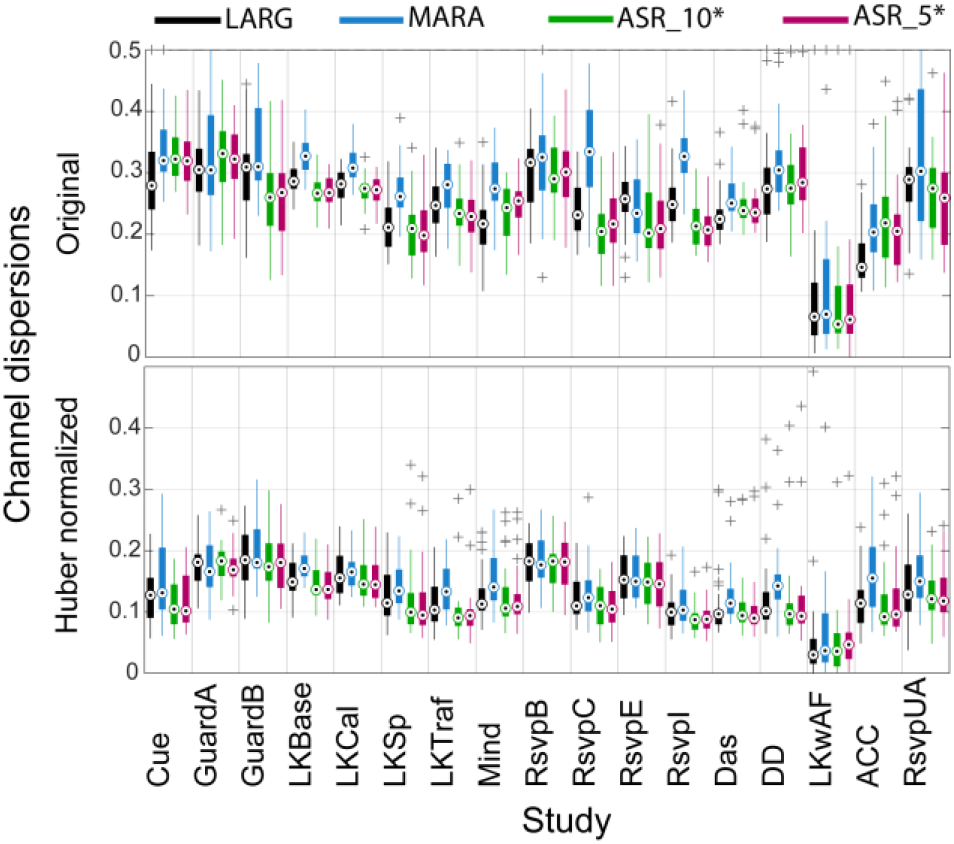
Channel dispersions by study before and after normalization by a recording-specific constant for four preprocessing methods: LARG (black), MARA (blue), ASR_10* (green), and ASR_5*(red).

### B. Effects of preprocessing on spectral characteristics

Fig. 3 summarizes the correlation between corresponding spectral features for various pairs of preprocessing methods based on correlations of corresponding spectral fingerprints.

**Fig. 3.**
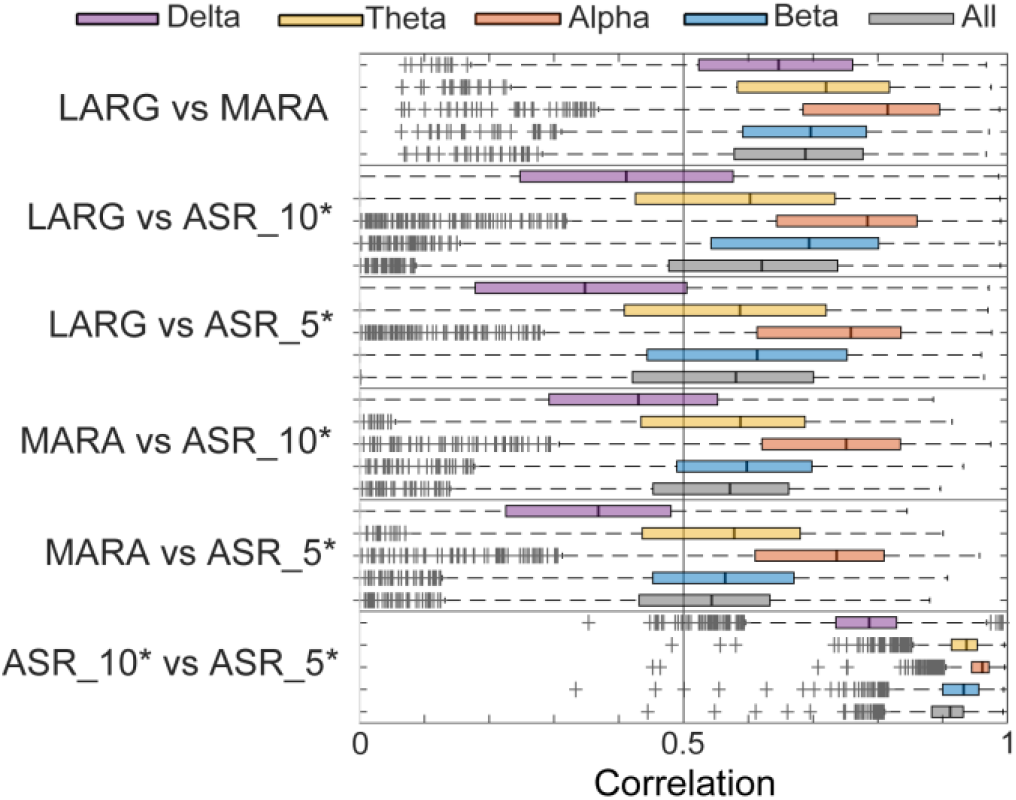
Evaluations of differences in signal spectral characteristics between pairs of preprocessing methods using spectral fingerprints.

As expected, the spectral samples of ASR_10* and ASR_5* are very highly correlated, and LARG and MARA have reasonably high spectral correlations. Even with these closely related pairs of methods, there are many outliers (appearing as a dark continuous bar due to the density of cross markers) with lower correlations. These low correlations most likely reflect differences in handling of artifacts between pipelines.

Disagreement between the ICA-based methods (LARG and MARA) and the ASR-based methods (ASR_10* and ASR_5*) in the delta frequency band ([2, 4] Hz for this analysis) is likely due to the differences in baselining and high-pass filtering that occurred at the beginning of the respective pipelines. However, ASR_10* and ASR_5* used the same input signals and even in this case, the correlations in the delta bands were much lower than in other bands. This suggests that not only should care be taken in specifying all baseline and preliminary filtering operations, but that small algorithmic differences in removal of large-amplitude low frequency artifacts such as blinks may affect downstream analysis in lower frequency bands. A comparison of spectral characteristics using the spectral sampling technique produces similar results. (See Fig S.2 in the supplementary material.)

### C. Relationships of event-related features across methods

Many EEG studies focus on event-related potentials (ERPs) in order to quantify the difference in evoked response due to an experimental factor, and it is important to ascertain whether any of these differences are due to variations in preprocessing methods. We looked at ERPs and ERSPs associated with different types of stimulus events for the 26 common channels across all 17 studies and calculated the correlations between corresponding features for different pairs of preprocessing methods as shown in Fig. 4.

**Fig. 4.**
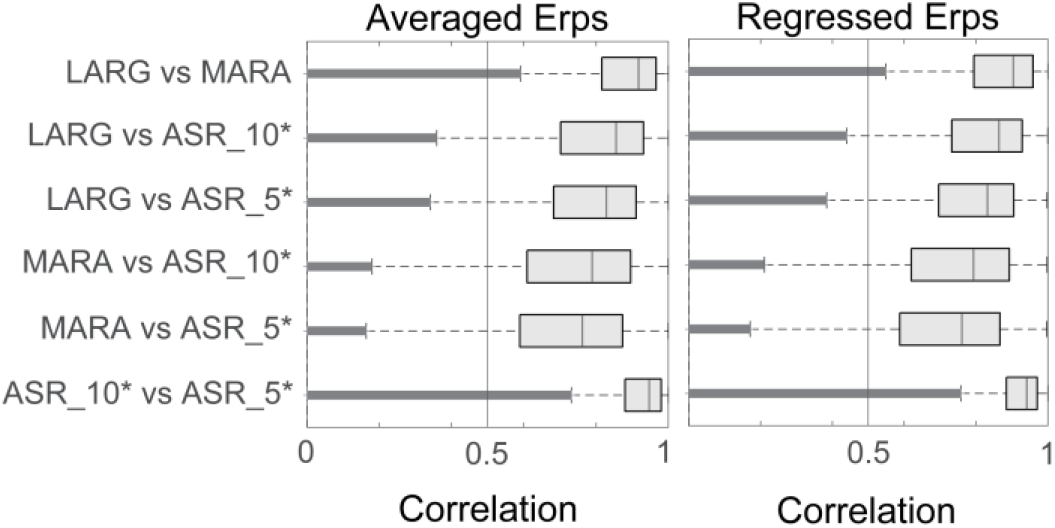
Correlations between corresponding event-related features produced by different pairs of preprocessing methods. Left boxplot shows ERPs and right boxplot shows ERSPs both computed by trial averaging. (Results using regression rather than trial averaging are similar and are not shown.)

Fig. 4 displays the distributions of correlations between corresponding features for pairs of preprocessing methods when ERPs (left graph) and ERSPs (right graph) are computed by trial averaging. Correlations are computed in the interval [0, 1] seconds. Fig. 4 uses boxplots to display the distributions of correlations between corresponding features for pairs of preprocessing methods when ERPs (left graph) and ERSPs (right graph) are computed by trial averaging. Correlations are computed in the interval [0, 1] seconds.

The graphs show that the relative levels of correlation between corresponding features are similar to those levels seen in the spectral analysis. The two variants of ASR are the most highly correlated although there are quite a few outlier features. LARG and MARA are more highly correlated for ERPs than either of those methods are with the ASR_5* and ASR_10*. LARG and ASR_10* are slightly more correlated than LARG and MARA for ERSPs.

For each pair of pre-processing methods, we used one-sample *t*-tests to test whether the mean of the distribution of ERP correlations (over all channels, analyzed events, and recordings) is significantly non-zero. Table 2 shows the means and 99% confidence intervals, confirming that the average correlation is significantly non-zero for all pairs of methods. Also shown in Table 2 are the median and the signed-rank statistic calculated using the Wilcoxon signed rank test for each pair of preprocessing methods. Regressed ERPs as well as averaged and regressed ERSPs gave similar statistical results. In all cases, the mean correlation was lower than the median.

**TABLE 2.** Correlations between corresponding averaged ERPs (DF = 840K)

| Preprocessing methods | $t$-test | | signed-rank test | |
| --- | --- | --- | --- | --- |
| | mean | conf int ($\alpha = 0.01$) | median | rstat |
| LARG $\nu$ MARA | 0.8588 | [0.8584, 0.8593] | 0.9175 | -184 |
| LARA $\nu$ ASR_10* | 0.7761 | [0.7755, 0.7768] | 0.8553 | -174 |
| LARG $\nu$ ASR_5* | 0.7627 | [0.7621, 0.7633] | 0.8286 | -152 |
| MARA $\nu$ ASR_10* | 0.7185 | [0.7179, 0.7192] | 0.7889 | -148 |
| MARA $\nu$ ASR_5* | 0.7017 | [0.7011, 0.7024] | 0.7618 | -131 |
| ASR_10* $\nu$ ASR_5* | 0.9037 | [0.9033, 0.9041] | 0.9469 | -175 |

To evaluate the consistency of features across recordings, for different preprocessing methods, we calculated the correlations among the features for recordings for each (study, event-code) triplet. We then performed paired *t*-tests and signed-rank tests both at the study and cross-study level to see which preprocessing methods produced the highest correlation for corresponding features across recordings. In all cases, both at the study level and at the corpus level, there was a strict statistically significant ordering of correlations: MARA > LARG > ASR_10* > ASR_5* with extremely small or vanishing *p*-values for both averaged and regressed features. That being said, the overall differences in correlations were very small. For regressed features, for example, the confidence intervals for the paired *t*-test comparisons were MARA−LARG: [0.001959, 0.002890], LARG−ASR_10*: [0.0095999, 0.010449], and ASR_10*−ASR_5*: [0.015595, 0.016365]. The statistics for averaged features were similar.

Although the feature correlations between preprocessing methods are similar, the actual features computed using trial averaging and regression have substantial differences. Fig. 5 displays study-wide feature averages for target events in three different RSVP studies for channel FCz. These studies were performed at three sites using three different Biosemi headsets. The top group of plots uses temporal overlap regression to compute regressed ERSPs (rERSPs), while the bottom group uses averaging (ERSPs). Outlier detection algorithms are incorporated in both averaging and regression techniques as described in [21]. Within a feature computation technique, agreement is fairly consistent and a prominent P300 apparent; although MARA appears to have removed most of this signal in the study averages of regressed features for RsvpB. ASR is known to have this issue for low burst cutoff thresholds, but for higher amplitude phenomena such as P300 there appears to be little difference between ASR_5* and ASR_10*. Importantly, the problematic nature of averaging is evident across all preprocessing methods. The bottom group of plots in Fig. 5 clearly shows the effect of other correlated and confounded events on ERSP estimation, with significant activity prior to the target event.

**Fig. 5.**
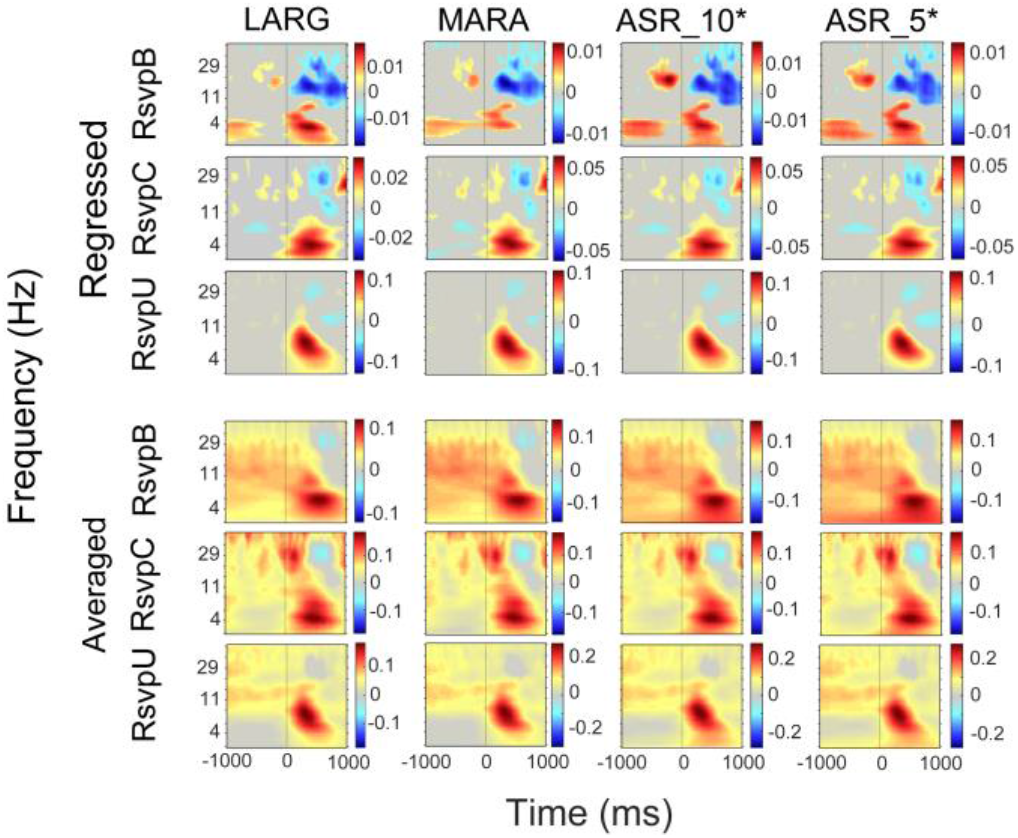
Comparison of study averages of (r)ERSP features for target events selected RSVP studies for channel FCz. Top group: averages of regressed ERSPs; bottom group: averaged ERSPs. Gray areas indicate lack of statistical significance (p > 0.01, FDR corrected).

### D. Effects of preprocessing on blink removal

Using box plots of the blink amplitude ratio, Fig. 6 summarizes how well the respective preprocessing methods remove blinks in the time domain.

**Fig. 6.**
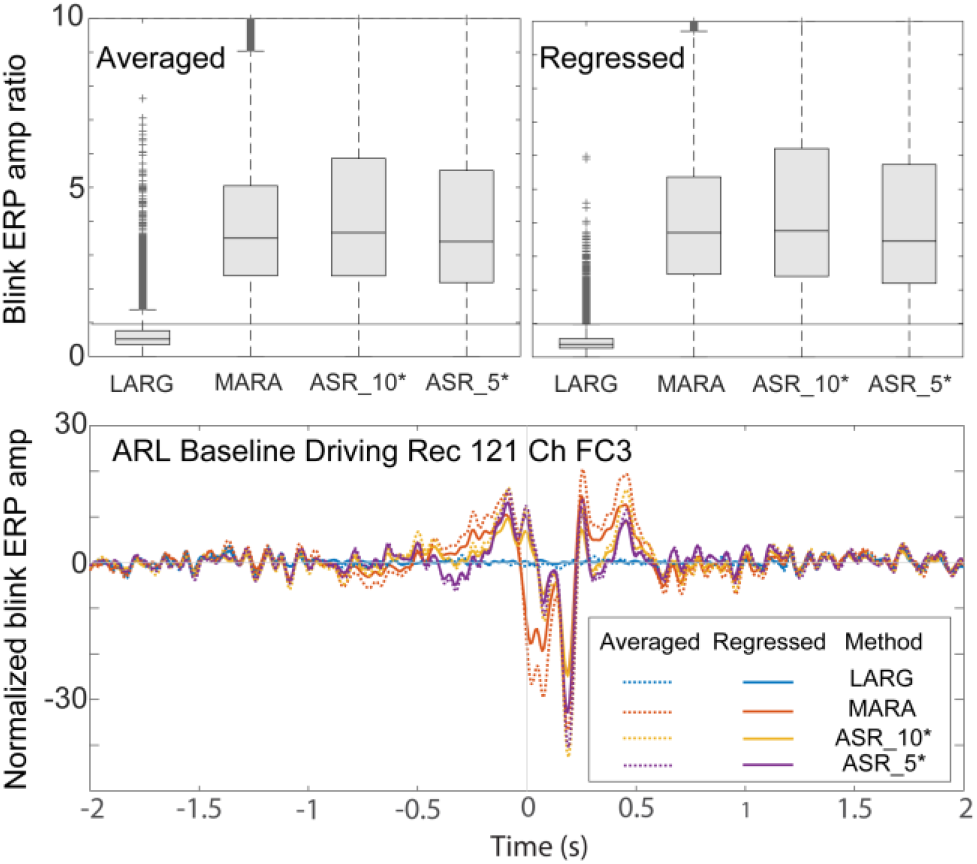
Temporal blink features after preprocessing. Top: Distribution of blink amplitude ratios for the corpus recordings using different preprocessing pipelines. Bottom: Blink (r)ERPs for a typical recording, with blink amplitude ratios closest to the median ratios. Blink amplitude ratios are computed for the 26 common channels in all recordings.

MARA and both variants of ASR display significant residuals in blink amplitude (ratio > 1) as shown by the extended whiskers in the corresponding box plots. In some recordings, this residual is very large, The ASR variations tend to leave more blink residual than MARA, while LARG tends to remove signal along with blinks (ratio <1). Signed-rank show a strict ordering of mean blink amplitude ratio of LARG << 1 << MARA < ASR_5* < ASR_10* with *p* values of essentially 0.

The bottom graph of Fig. 6 shows a typical blink ERPs overlaid for different preprocessing methods and different computation strategies. The ERP versions have been scaled by subtracting the mean in the intervals [−2, −1.5] and [1.5, 2] and then dividing by the median absolute value of the resulting amplitude in those subintervals. The particular recording whose (r)ERPs were chosen is the one whose blink amplitude ratio was closest to the individual median blink amplitude ratios for the different preprocessing methods.

This example is typical of the others that we have examined. The residual signal is quite large for all preprocessing methods except LARG, which directly regresses out the blink signal in the interval [−1, 1]. In this example (which is typical), the other methods appear to remove too much signal at the blink maximum and too little signal before and after the maximum. The averaged and regressed blink ERPs are close for the ASR variants, but the averaged blink ERP for MARA shows more blink residual than its regressed version.

All four preprocessing methods show spectral blink residuals. Fig. 7 compares the study averages of the rERSPs associated with the blink maximum event for three different studies. GuardA is a complex, time-extended visual search task, LKCal is a simulated vehicular lane-keeping task, while RSVPI is a demanding, time-compressed visual target detection task.

**Fig. 7.**
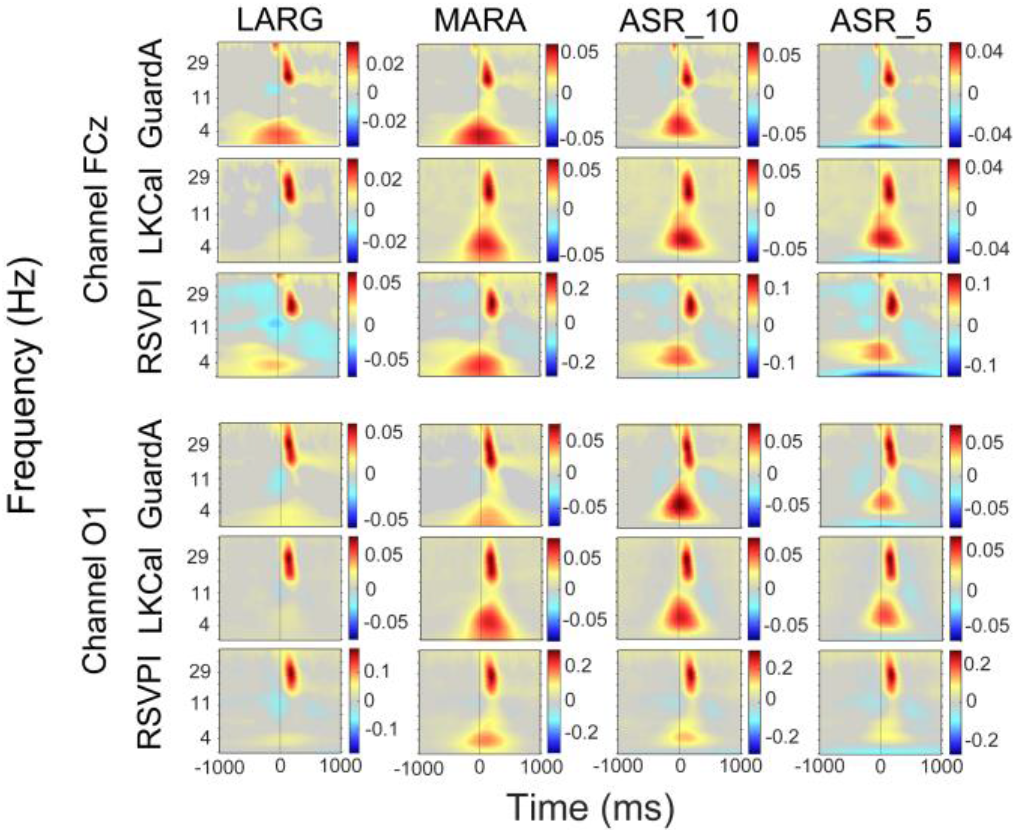
Comparison of regressed ERSPs of blink events for 3 selected studies. The top group displays the results for channel FCz and the bottom group displays the results for channel O1. Gray areas indicate lack of statistical significance (*p* > 0.01, FDR corrected).

The top group shows channel FCz, while the bottom group shows channel O1. All of the methods exhibit a significant burst-like increase in power in the beta frequency range occurring slightly after the blink maximum, possibly associated with the beginning of the eye opening phase. MARA, and to a lesser extent the ASR variants, show significant low-frequency activity time-locked to the blink maximum, which could be associated with residual blink activity.

## IV. Discussion

This paper investigated differences in outcome at various stages in analysis due to choices made during processing. We focused on two types of processing approaches: ICA-based (LARG and MARA) and subspace reconstruction (ASR_5* and ASR_10*). Our large-scale analysis shows that the resulting signals have generally similar characteristics, but there are small systematic differences in outcomes, even between closely related methods.

### A. Eye artifacts affect signal characteristics

The characteristics of signals with just external artifacts removed (top row of Fig. 1) are dramatically different than the characteristics of signals in which subject-generated artifacts (rows 2 through 5 of Fig. 1) are also removed. Fig. 1 also shows that the global signal characteristics after subject-generated artifacts are removed are very similar across preprocessing methods. Since LARG mainly focuses on the removal of blinks and eye artifacts, one can conclude, as expected, that the majority of the large-scale signal differences are due to blinks.

These methods produce data in which blinks are difficult to observe in single trials. LARG, which directly regresses out blinks during preprocessing, has blink amplitude ratios less than 1, leading to the concern that perhaps too much EEG signal has been removed, while the other preprocessing methods may not remove enough of the blink artifacts (Fig. 6). All of the methods, including LARG, show similar well-defined time-frequency features time-locked to blink events after blink removal (Fig. 7).

We also computed (not shown here) blink amplitude ratios of epochs time-locked to the blink maximum for selected individual recordings. The blink ratios for the individual epochs are very close to 1 for all methods and indistinguishable from randomly selected epochs that contain no blinks. However, when we average these epochs for a single recording, the blink ratios are in agreement with those reported in Fig. 6. This suggests that direct viewing of the signals after preprocessing, may lead to a false conclusion that the blink effects have been mostly removed, when in fact there are systematic biases in blink epochs. Blum et al. [22] recently compared blink removal in the regular ASR algorithm and a modification based on Riemannian geometry. They also observed systematic blink residuals that may be indicative of event-related potentials associated with eye blinks.

Blink entrainment in certain visual tasks can further complicate the interpretation [23]. We recommend that researchers generally assume residual blink signals are present in their data after preprocessing and take active measures to address this when interpreting their results. However, researchers have observed neural activity locked to spontaneous blinks. This is hypothesized to be related to attentional disengagement and transient activation/deactivation of cortical brain networks [24]. It is therefore important to examine multiple factors, including the spatial and spectral distribution of residual activity locked to blinks, when characterizing the origin of this activity. Temporal overlap regression [5] [6] may also be a particularly suitable method to address this problem by regressing out common patterns of activity unique to blink events.

### B. Scaling by a constant reduces inter-recording variability

As we reported in earlier work [19], our results highlight the potential for factoring out a portion of inter-recording variability by uniform scaling of channel amplitudes (Fig. 2). This simple step is effective across preprocessing methods and is strongly recommended for cross-recording comparisons, even within a single study. This scaling does not change the relative sizes of the respective channel amplitudes.

### C. Filtering and pre-processing differences

Filtering and its effects on EEG signals is a complex issue that has been examined by a number of authors [25]. Widmann et al. [26] provide useful guidelines, pointing out that filter design trade-offs are highly dependent on the nature of the problem being addressed and on the signal quality. In this paper, we opted for high-pass filtering using FIR non-casual zero-phase filtering with Hamming windows for all preprocessing methods. We recognize that this choice is limiting for certain applications, and that a large-scale study of signal distortion for different filtering alternatives would be useful. Universal recommendations for filter selection are probably not possible, even if trade-offs are well-documented.

One place where there was a distinct difference in the choice of filter parameters was in the high-pass filter used for ASR versus the other preprocessing methods. ASR depends on the signal having zero mean, both globally, as well as over local (e.g., 0.5 sec) analysis windows. High-pass filtering is an effective way to remove local signal drift and produce a zero-mean time series. However, EEG recording hardware from some manufacturers, such as Biosemi, have large DC offsets or drift that may require a suitably large stop-band suppression in the filter to ensure that power at 0 Hz (corresponding to the mean) is as close to zero as possible. In this work, we used an FIR high-pass filter with a 0.75-1.5 Hz transition band to achieve 70 dB reduction in power at 0.5 Hz using the same FIR filtering approach used for LARG and MARA. However, since LARG and MARA used a 0.5-1 Hz transition band, we cannot rule out that some differences observed between these methods and ASR, particularly in the delta band, may be attributed to differences in the filter cutoff. However, since each pair of methods (pair 1: LARG and MARA; pair 2: ASR_10* and ASR_5*) used the same input within the pair, the large spectral differences within each pair of methods is likely attributable to differences in artifact handling not filtering (Fig. 3).

Another difference between the input signals to the four preprocessing pipelines is that ASR requires full-rank data and thus cannot be applied after channel interpolation, ICA component removal, or other rank-reducing methods. However, the comparison metrics described here require a fixed, common set of channels. LARG and MARA interpolated bad channels prior to performing their analysis and used PCA to reduce rank. The normal offline ASR algorithm operates after bad channels have been removed. ASR* just dealt with the bad channels as part of its subspace removal and did relatively well. The effects of channel interpolation should be further investigated.

### D. Event-related features

ERPs have been used in restricted experimental settings to assess processing or headsets effects. Barham et al. [27] compared correlations of individual target and non-target trials for 15 subjects in an auditory oddball task. They also compared the N200 and P300 amplitudes and latencies between standard and deviate trials. Cruz-Garza et al. [20] used a spectral clustering approach to quantify headset differences since direct comparison was not possible across different datasets.

Fig. 6 shows that, although one might expect roughly similar event-related features across preprocessing methods, the details of individual corresponding features may differ considerably. Even the two ASR variants, which have a median feature correlation greater than 0.9, have many outlier examples with very low correlation. All of the event-related features computed in this paper used a trial outlier method that excludes epochs with unusually large amplitudes. Identifying other types of artifactual trials before preprocessing and systematically examining how excluding these trials changes the feature, may be useful in evaluating feature generalizability.

To improve the generalizability, we uses a diverse set of stimulus events and extended the comparison to event-locked time-frequency features and regression-derived features. Our conclusions are generally consistent across feature and event types. Fig. 5 shows that ERSPs computed by trial averaging can have significant mixing of evoked responses from temporally adjacent events, particularly for RSVP paradigms that elicit overlapping activity from rapidly presented stimuli.

## V. Conclusion

In this work, we characterized the effects of four pre-processing approaches on spatial, spectral, and temporal EEG features across 17 heterogeneous studies. Our results suggest that even small changes in artifact removal strategy may result in differences with large effects on particular portions of the signal. While there is general agreement on the steps that should be taken for preprocessing (e.g., filtering, line-noise removal, references, bad channels handling, artifact removal), a range of “standard” choices may affect results in unknown ways. While differences may be small when averaged over a large, diverse corpus, they may be significant when considered for a single study. Rather than anoint a particular analysis path as the “gold standard”, a diversity approach may lead to more reproducible and meaningful results. If a federation of automated processing pipelines with well-documented parameter choices were available, researchers could run their data through several of them and compare the results as part of reporting their research. Large differences in analysis output would be analyzed as part of the research reporting, leading to a better understanding both of the methods and the underlying neural phenomena. In addition to using regression instead of averaging to calculate event-related features, we also recommend that researchers analyze the distribution of blinks relative to other events.

## Acknowledgment

The authors would like to thank Tony Johnson and Michael Dunkel for data assembly, and the experimenters, including Ching-Teng Lin and Jung-Tai King of NCTU, who contributed their data. This work received computational support from UTSA’s HPC cluster Shamu. Research was sponsored by the United States Army Research Laboratory under Cooperative Agreement Number W911NF-10-2-0022. The views and the conclusions contained in this document are those of the authors and should not be interpreted as representing the official policies, either expressed or implied, of the Army Research Laboratory or the U.S Government. The U.S Government is authorized to reproduce and distribute reprints for Government purposes notwithstanding any copyright notation herein.

---

**Supplemental figure 1:**
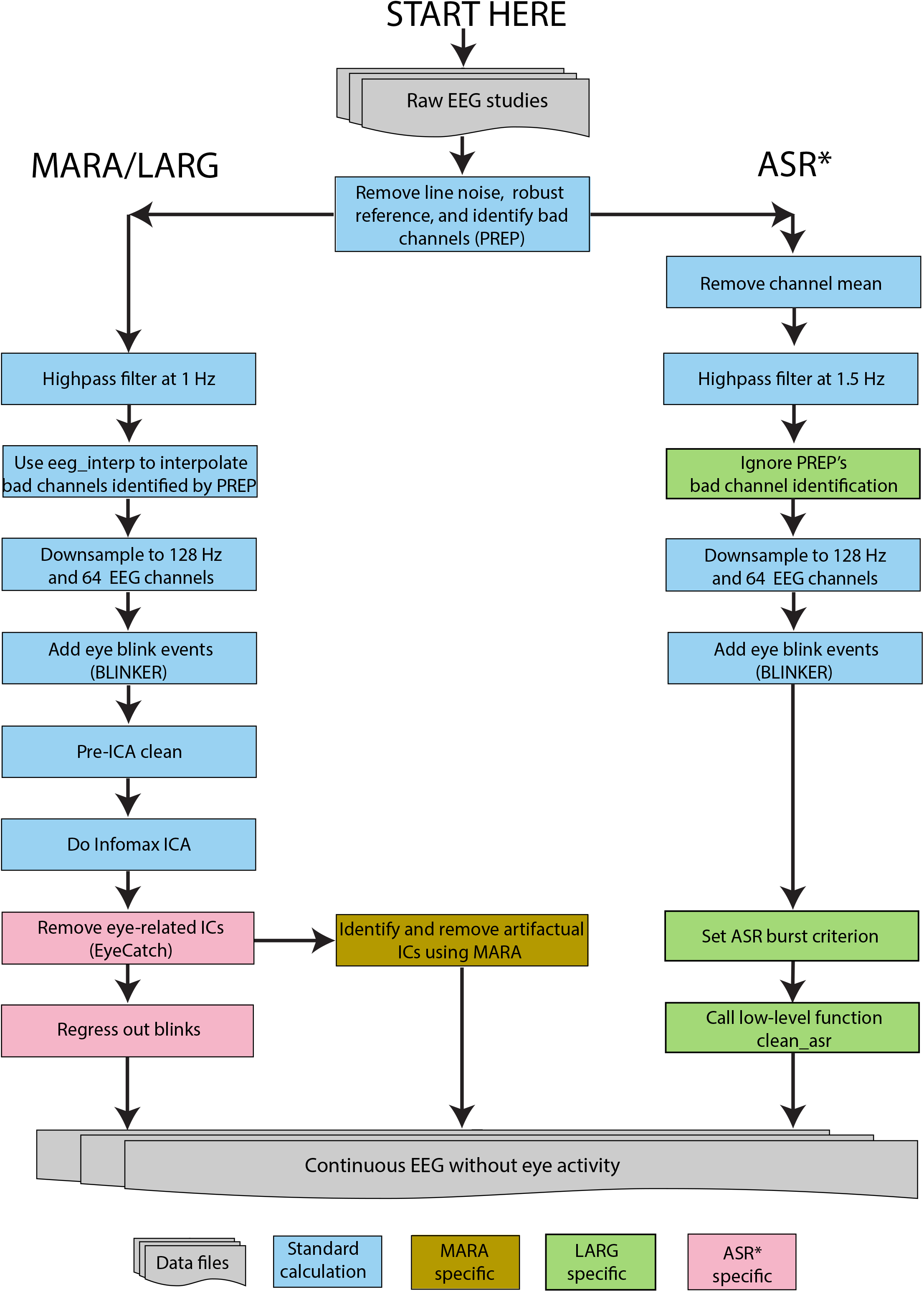
Summary of pipelines

**Supplemental Figure 2:**
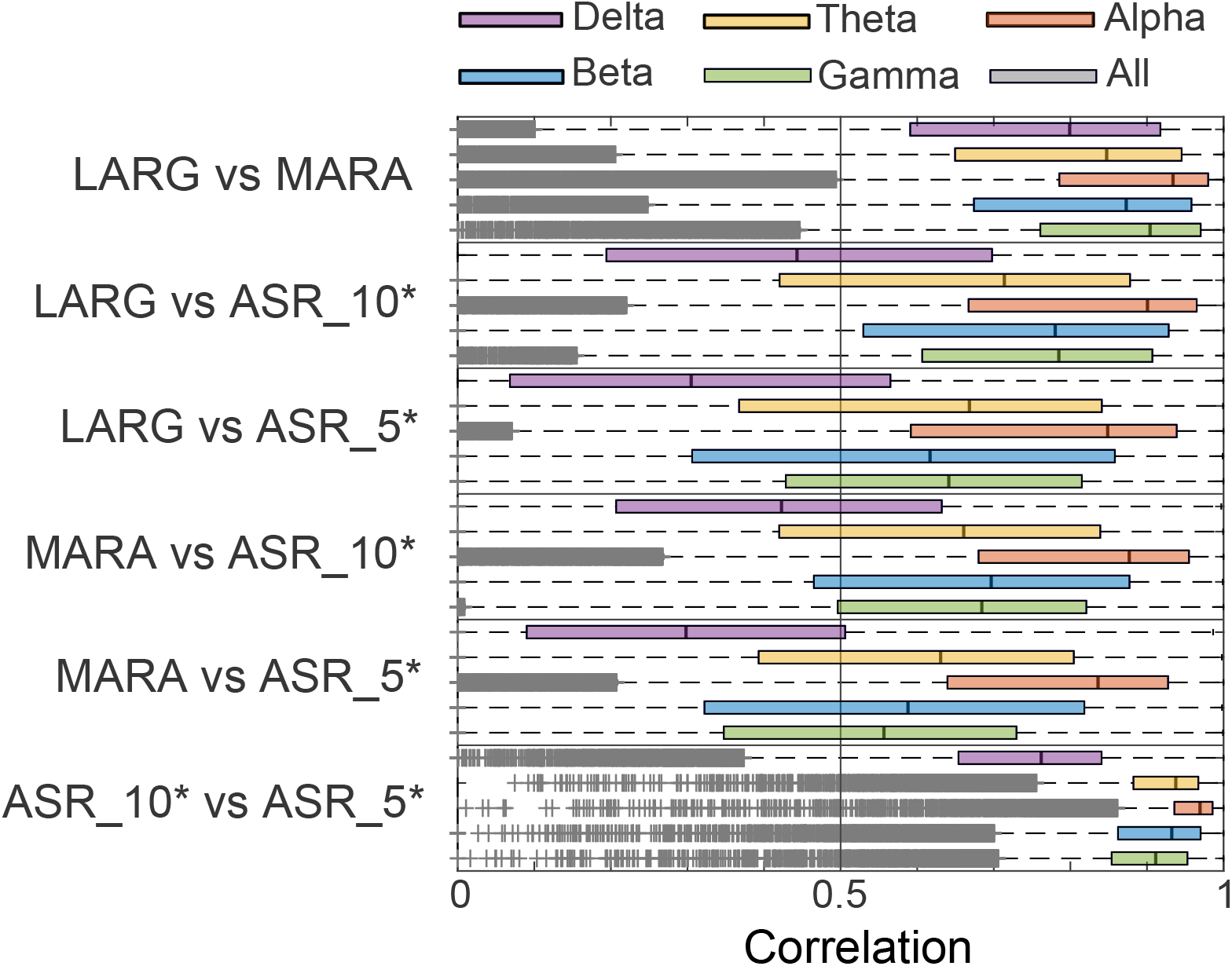
Random sample spectral correlations

## References

[1] J. Gross et al., “Good practice for conducting and reporting MEG research,” NeuroImage, vol. 65, pp. 349–363, Jan. 2013, doi: 10.1016/j.neuroimage.2012.10.001.

[2] A. Keil et al., “Committee report: Publication guidelines and recommendations for studies using electroencephalography and magnetoencephalography,” Psychophysiology, vol. 51, no. 1, pp. 1–21, 2014, doi: 10.1111/psyp.12147.

[3] C. R. Pernet et al., “Best practices in data analysis and sharing in neuroimaging using MEEG,” 2018, doi: 10.31219/osf.io/a8dhx.

[4] V. Litvak, A. Jha, G. Flandin, and K. Friston, “Convolution models for induced electromagnetic responses,” NeuroImage, vol. 64, pp. 388–398, Jan. 2013, doi: http://dx.doi.org.libweb.lib.utsa.edu/10.1016/j.neuroimage.2012.09.014.

[5] E. Kristensen, A. Guerin-Dugué, and B. Rivet, “Regularization and a general linear model for event-related potential estimation,” Behav. Res. Methods, pp. 1–20, Mar. 2017, doi: 10.3758/s13428-017-0856-z.

[6] B. V. Ehinger and O. Dimigen, “Unfold: an integrated toolbox for overlap correction, non-linear modeling, and regression-based EEG analysis,” PeerJ, vol. 7, p. e7838, Oct. 2019, doi: 10.7717/peerj.7838.

[7] N. Bigdely-Shamlo, J. Touryan, A. Ojeda, C. Kothe, T. Mullen, and K. Robbins, “Automated EEG mega-analysis I: Spectral and amplitude characteristics across studies,” NeuroImage, p. 116361, Nov. 2019, doi: 10.1016/j.neuroimage.2019.116361.

[8] N. Bigdely-Shamlo et al., “Hierarchical Event Descriptors (HED): Semi-structured tagging for real-world events in large-scale EEG,” Front. Neuroinformatics, vol. 10, 2016, doi: 10.3389/fninf.2016.00042.

[9] N. Bigdely-Shamlo, T. Mullen, C. Kothe, K.-M. Su, and K. A. Robbins, “The PREP pipeline: standardized preprocessing for large-scale EEG analysis,” Front. Neuroinformatics, vol. 9, 2015, doi: 10.3389/fninf.2015.00016.

[10] K. Kleifges, N. Bigdely-Shamlo, S. E. Kerick, and K. A. Robbins, “BLINKER: Automated extraction of ocular indices from EEG enabling large-scale analysis,” Front. Neurosci., vol. 11, 2017, doi: 10.3389/fnins.2017.00012.

[11] I. Winkler, S. Haufe, and M. Tangermann, “Automatic classification of artifactual ICA-components for artifact Removal in EEG signals,” Behav. Brain Funct., vol. 7, p. 30, 2011, doi: 10.1186/1744-9081-7-30.

[12] C. A. E. Kothe and T.-P. Jung, “Artifact removal techniques with signal reconstruction,” WO2015047462A9, 16-Jul-2015.

[13] A. Delorme and S. Makeig, “EEGLAB: an open source toolbox for analysis of single-trial EEG dynamics including independent component analysis,” J. Neurosci. Methods, vol. 134, no. 1, pp. 9–21, Mar. 2004, doi: 10.1016/j.jneumeth.2003.10.009.

[14] A. Delorme et al., “EEGLAB, SIFT, NFT, BCILAB, and ERICA: New tools for advanced EEG processing,” Computational Intelligence and Neuroscience, 2011. [Online]. Available: https://www.hindawi.com/journals/cin/2011/130714/. [Accessed: 14-Aug-2017].

[15] T. R. Mullen et al., “Real-time neuroimaging and cognitive monitoring using wearable dry EEG,” IEEE Trans. Biomed. Eng., vol. 62, no. 11, pp. 2553–2567, Nov. 2015, doi: 10.1109/TBME.2015.2481482.

[16] F. Raimondo, J. E. Kamienkowski, M. Sigman, and D. Fernandez Slezak, “CUDAICA: GPU optimization of Infomax-ICA EEG analysis,” Computational Intelligence and Neuroscience, 2012. [Online]. Available: https://www.hindawi.com/journals/cin/2012/206972/. [Accessed: 20-Apr-2019].

[17] N. Bigdely-Shamlo, K. Kreutz-Delgado, C. Kothe, and S. Makeig, “EyeCatch: Data-mining over half a million EEG independent components to construct a fully-automated eye-component detector,” in 2013 35th Annual International Conference of the IEEE Engineering in Medicine and Biology Society (EMBC), 2013, pp. 5845–5848, doi: 10.1109/EMBC.2013.6610881.

[18] I. Winkler, S. Brandl, F. Horn, E. Waldburger, C. Allefeld, and M. Tangermann, “Robust artifactual independent component classification for BCI practitioners,” J. Neural Eng., vol. 11, no. 3, p. 035013, 2014, doi: 10.1088/1741-2560/11/3/035013.

[19] N. Bigdely-Shamlo, J. Touryan, A. Ojeda, C. Kothe, T. Mullen, and K. Robbins, “Automated EEG mega-analysis I: Spectral and amplitude characteristics across studies,” NeuroImage, p. 116361, Nov. 2019, doi: 10.1016/j.neuroimage.2019.116361.

[20] J. G. Cruz-Garza et al., “Deployment of mobile EEG technology in an art museum setting: Evaluation of signal quality and usability,” Front. Hum. Neurosci., vol. 11, 2017, doi: 10.3389/fnhum.2017.00527.

[21] N. Bigdely-Shamlo, J. Touryan, A. Ojeda, C. Kothe, T. Mullen, and K. Robbins, “Automated EEG mega-analysis II: Cognitive aspects of event related features,” NeuroImage, p. 116054, Sep. 2019, doi: 10.1016/j.neuroimage.2019.116054.

[22] S. Blum, N. S. J. Jacobsen, M. G. Bleichner, and S. Debener, “A Riemannian nodification of Artifact Subspace Reconstruction for EEG artifact handling,” Front. Hum. Neurosci., vol. 13, 2019, doi: 10.3389/fnhum.2019.00141.

[23] C. Kranczioch, “Individual differences in dual-target RSVP task performance relate to entrainment but not to individual alpha frequency,” PLOS ONE, vol. 12, no. 6, p. e0178934, Jun. 2017, doi: 10.1371/journal.pone.0178934.

[24] M. E. Raichle, A. M. MacLeod, A. Z. Snyder, W. J. Powers, D. A. Gusnard, and G. L. Shulman, “A default mode of brain function,” Proc. Natl. Acad. Sci. U. S. A., vol. 98, no. 2, pp. 676–682, Jan. 2001, doi: 10.1073/pnas.98.2.676.

[25] G. A. Rousselet, “Does filtering preclude us from studying erp time-courses?,” Front. Psychol., vol. 3, May 2012, doi: 10.3389/fpsyg.2012.00131.

[26] A. Widmann, E. Schröger, and B. Maess, “Digital filter design for electrophysiological data – a practical approach,” J. Neurosci. Methods, vol. 250, pp. 34–46, Jul. 2015, doi: 10.1016/j.jneumeth.2014.08.002.

[27] M. P. Barham, G. M. Clark, M. J. Hayden, P. G. Enticott, R. Conduit, and J. A. G. Lum, “Acquiring research-grade ERPs on a shoestring budget: A comparison of a modified Emotiv and commercial SynAmps EEG system,” Psychophysiology, vol. 54, no. 9, pp. 1393–1404, 2017, doi: 10.1111/psyp.12888.

